# TGF-βRII knock-down promotes tumor growth and chemoresistance to gemcitabine of pancreatic cancer cells via phosphorylation of STAT3

**DOI:** 10.1101/352963

**Authors:** Vincent Drubay, Nicolas Skrypek, Lucie Cordiez, Romain Vasseur, Céline Schulz, Nihad Boukrout, Bélinda Duchêne, Lucie Coppin, Isabelle Van Seuningen, Nicolas Jonckheere

## Abstract

Pancreatic adenocarcinoma (PDAC) is one of the most deadly cancers in the western countries because of a lack of early diagnostic markers and efficient therapeutics. At the time of diagnosis, more than 80% of patients have metastasis or locally advanced cancer and are therefore not eligible for surgical resection. Pancreatic cancer cell also harbour a high resistance to chemotherapeutic drugs such as gemcitabine that is one of the main palliative treatment for PDAC.

TGF-β possesses both tumor-suppressive and oncogenic activities in pancreatic cancer. TGF-β signalling pathway plays complex role during carcinogenesis by initially inhibiting epithelial growth and later promoting the progression of advanced tumors and thus emerged as tumor suppressor pathway. TGF-β binds to its receptor TGF-βRII and activates different pathways: canonical pathway involving the Smad proteins and alternative pathways such as MAPKs. Smad4 is mutated in 50-80% of PDAC. Mutations of TGF-βRII also occurs (5-10%). In order to decipher the role of TGF-β in carcinogenesis and chemoresistance, we decided to characterize the knocking down of TGF-βRII that is the first actor of TGF-β signalling. We developed pancreatic cancer cell lines stably invalidated for TGF-βRII and studied the impact on biological properties of pancreatic cancer cells both in vitro and in vivo. We show that TGF-βRII silencing alters tumor growth and migration as well as resistance to. TGF-βRII silencing also leads to S727 STAT3 and S-63 c-Jun phosphorylation, decrease of MRP3 and increase of MRP4 ABC transporter expression and induction of a partial EMT phenotype.

In the future, the better understanding TGF-β signaling pathways and underlying cellular mechanisms in chemoresistance to gemcitabine may bring new therapeutic tools to clinicians.

## Introduction

Pancreatic cancers (PC) are projected to become the second leading cause of cancer-related death by 2030 (Rahib et al., 2014). The survival curve is extremely short (6 months) and the survival rate at 5 years is very low (3%). This dramatic outcome is related to a lack of therapeutic tools and early diagnostic markers which makes pancreatic cancer the most deadly cancer. At the time of diagnosis, more than 80% of PC are already metastatic or locally advanced and only about 10 to 15% of patients are considered eligible for surgical resection (Vincent et al., 2011). Remaining patients that do not benefit of surgery will receive palliative chemotherapy and notably gemcitabine, a fluorinated analog of deoxycytidine that is a major chemotherapeutic drug used in firstline in advanced PC. Unfortunately, PC is characterized by an intrinsic and acquired chemoresistance that lead to relapse and death (Kleeff et al., 2016). Deciphering mechanisms responsible for PC cell resistance to gemcitabine is thus crucial to improve efficacy of the drug and propose more efficient therapies.

TGF-β signalling pathway plays a complex role during carcinogenesis. TGF-β initially inhibits epithelial growth whereas it appears to promote the progression of advanced tumors and thus emerged as tumor suppressor pathway in pancreatic cancer (Principe et al., 2014). TGF-β can act in an autocrine manner or as a paracrine factor secreted by the microenvironment (Derynck et al., 2001). After binding to its receptor TGF-βRII, TGF-β signals via activation of several pathways. The canonical pathway involves the Smad proteins, but activation of other pathways such as MAPKs, PI3K or small GTPases (Derynck et al., 2001) pathways may also mediate TGF-β effects. It is also interesting to note that Smad4/DPC4 (deleted in pancreatic cancer 4) is mutated in 50-80% of PDAC whereas mutations of TGF-βRII are less common (5-10%) (Kleeff et al., 2016; TCGA-Network., 2017). We previously showed that TGF-β can regulate gene expression via canonical or alternative signalling pathways (Jonckheere et al., 2004).

In order to design new therapeutic strategies, it is thus mandatory to better characterize the signaling pathways and complex gene networks that are altered during carcinogenesis progression. Therefore, to better understand the role and contribution of TGF-βRII in TGF-β signalling and biological properties of PC cells in vitro and in vivo, we developed PC cell lines stably invalidated for TGF-βRII. Our results show that TGF-βRII silencing alters tumor growth and migration and increases resistance to gemcitabine in vitro and in vivo. TGF-βRII silencing also leads to STAT3 and c-Jun phosphorylation, alteration of MRP3 and MRP4 ABC transporters expression and induction of a partial EMT phenotype.

This work underlies the importance of TGF-β signaling pathways and associated cellular mechanisms as inducers of chemoresistance to gemcitabine and proposes potential new therapeutic tools to clinicians, surgeons and anatomopathologists for this deadly disease.

## Material and methods

### Cell culture

CAPAN-1 and CAPAN-2 PC cell lines were cultured as previously described (Jonckheere et al., 2004). TGF-βRII-knocked down (KD) cells were obtained following stable transfection of CAPAN-1 and CAPAN-2 cells with four different pGeneClipTM puromycin vectors encoding TGF-βRII ShRNA (SA BiosciencesTM) as previously described (Jonckheere et al., 2012). The empty vector was used to raise control clones called Non Targeting (NT). Four selected clones of NT and each TGF-βRII-KD cells were pooled in order to avoid clonal variation and were designated TGF-βRIIKD6, TGF-βRIIKD7, TGF-βRIIKD8 and TGF-βRIIKD9. All cells were maintained in a 37°C incubator with 5% CO2 and cultured as the parental cells.

### qRT-PCR

Total RNA from PC cells was prepared using the NucleoSpin^®^ RNA II kit (Macherey Nagel, Hoerdt, Germany). cDNA was prepared as previously described (Van Seuningen et al., 2000). Semi-quantitative PCR was performed as previously described (Mesquita et al., 2003). qPCR was performed using SsoFastTM Evagreen Supermix kit following the manufacturer’s protocol using the CFX96 real time PCR system (Bio-Rad). Primer information is given in table 1. Each marker was assayed in triplicate in three independent experiments. Expression level of genes of interest was normalized to the mRNA level of GAPDH housekeeping gene.

**Table 1.**
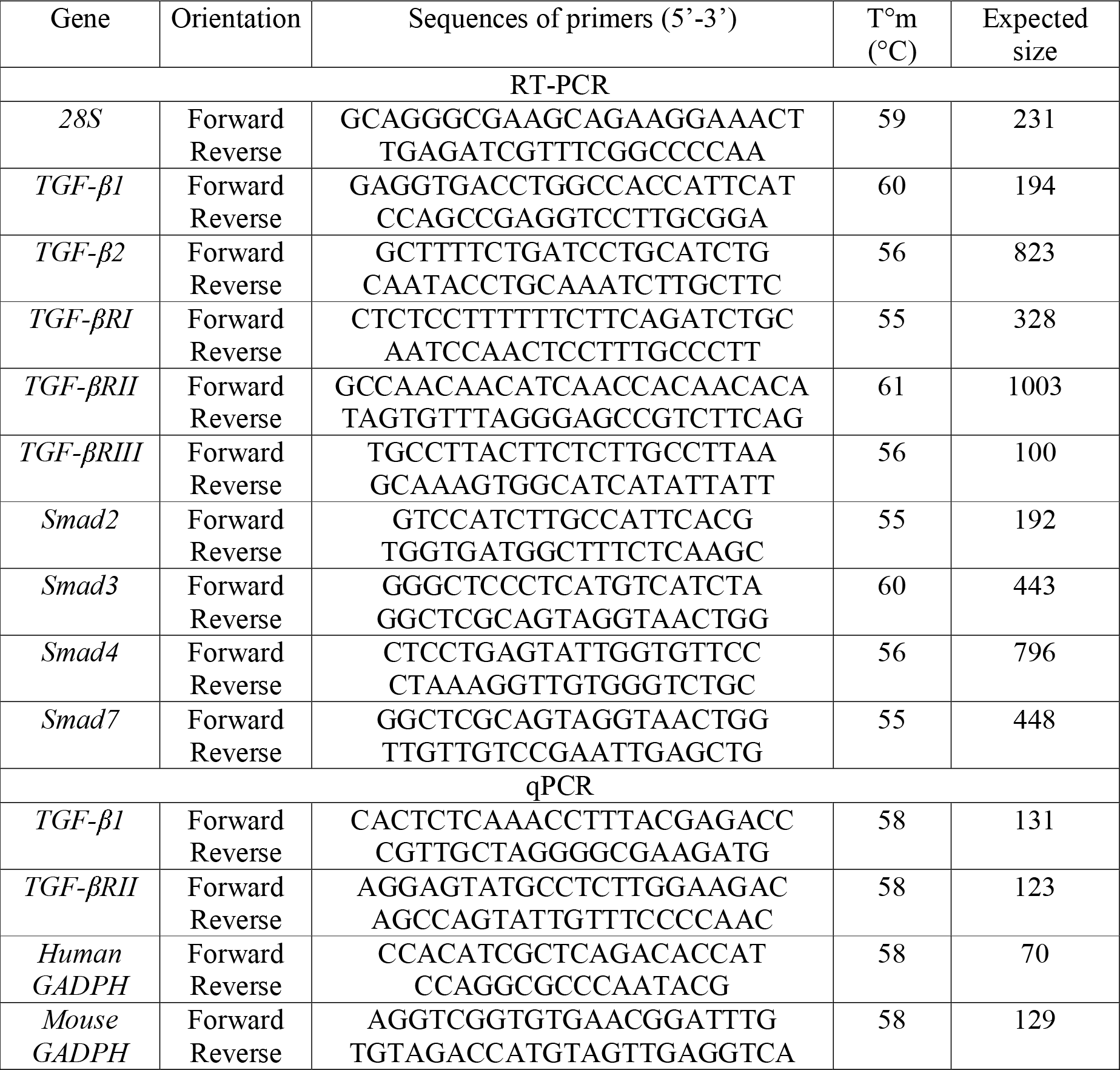
Primers used for RT-PCR and qPCR experiments.

### Protein extraction and western-blotting

Total cellular extracts were performed as previously described in Van Seuningen et al. (Van Seuningen et al., 1995) and Jonckheere et al. (Jonckheere et al., 2009). Western-blotting on nitrocellulose membrane (0.2 μm, Whatman) was carried out as previously described (Piessen et al., 2007). Membranes were incubated with antibodies against STAT3 (79D7, Cell signalling), phospho S727 STAT3 (9134, signalling), c-Jun (60A8, Cell signalling), phospho S63 c-Jun (54B3, Cell signalling) and β-actin (AC-15, sigma). Antibodies were diluted in 5% (w/v) non-fat dry milk in Tris-Buffered Saline Tween-20 (TBS-T). Peroxydase-conjugated secondary antibodies (Sigma-Aldrich) were used and immunoreactive bands were visualised using the West Pico chemoluminescent substrate (Thermo Scientific, Pierce, Brebières, France). Chemo-luminescence was visualised using LAS4000 apparatus (Fujifilm). Density of bands were integrated using Gel analyst software^®^ (Claravision, Paris, France) and represented as histograms. Three independent experiments were performed.

### Cell proliferation

Cells were seeded at 1×10^5^ cells per well in 6-well plates. Cells were counted daily using a Malassez counting chamber using Trypan Blue exclusion dye (Life Technologies) during 96h. Experiments were performed three times in triplicate.

### Wound healing test

1500 cells were seeded per wells in 96 well plates (Image LockTM plates, Essen Bioscience) and cultured until confluence was reached. The wound was realized using IncuCyte wound maker (Essen BioScience). Cells were washed three times with PBS 1X and complete medium was added to the cells. Wound widths were analyzed using Incucyte platform (Live-Cell imaging System, Essen Bioscience) and pictures collected every 2h.

### Cytotoxicity assay

Cells were seeded in growth medium into 96-well plates at a density of 10^4^ cells per well. After 24h incubation, the medium was replaced by fresh medium containing gemcitabine at 35 nM and incubated for 72h at 37°C. The viability of cells was determined using the 3-(4,5-dimethylthiazol-2-yl)-2,5-diphenyltetrazolium bromide assay (MTT, Sigma-Aldrich) as previously described (Skrypek et al., 2013). Percentage of viability = [(A_treated_ – A_blan_k)/(A_neg._ – A_blank_)] × 100; where A_treated_ is the average of absorbance in wells containing cells treated with gemcitabine, A_neg._ is the average of wells containing cells without gemcitabine treatment, and A_blank_ is the average of wells containing medium without cells.

### Subcutaneous xenografts

NT or TGF-βRII-KD CAPAN-1 (10^6^ cells in 100 μl Matrigel) and CAPAN-2 (2×10^6^) cells were injected subcutaneously (SC) into the flank of seven-week-old male Severe Combined Immunodeficient (SCID) mice (CB17, Janvier, France). Six mice were used per group. Tumor size was evaluated weekly by measuring the length (l) and the width (L) and tumor volume was calculated with the formula (I^2^×L). Once palpable tumors were developed (250 mm^3^), gemcitabine (15 mg/kg) or PBS (200 μl) were injected intra-peritoneously, twice a week. All procedures were in accordance with the guideline of animal care committee (Comité Ethique Expérimentation Animale Nord Pas-de-Calais, #122012).

### Immunohistochemistry

Xenografts were fixed in 10% (w/v) buffered formaldehyde, embedded in paraffin, cut at 4 μm thickness and applied on SuperFrost^®^ slides (Menzel-Glaser, Braunschweig, Germany). Manual IHC was carried out as previously described (van der Sluis et al., 2004). The antibodies were used as followed: anti-STAT3 (1:200, #483 Santa Cruz), anti-c-Jun (1:200, 60A8 Cell signaling), anti-E-Cadherin (1:200, 3195 Cell signalling) and anti-vimentin (1:200, sc5741, Santa Cruz). Intensity of staining was graded as weak (1), moderate (2) or strong (3). The percentage of ductal stained cells was graded as 1 (0-25%), 2 (25-50%), 3 (50-75%) and 4 (75-100%). Total score was calculated by multiplying the intensity score and percentage score.

### Expression analysis in CCLE database

TGF-βRII, ABCB1/MDR1, ABCC1/2/3/4/5 and ABCG2 z-score expressions were extracted from databases available at cBioPortal for Cancer Genomics (Cerami et al., 2012; Gao et al., 2013). The queries were realized in CCLE (44 pancreatic samples, Broad Institute, Novartis Institutes for Biomedical Research) (Barretina et al., 2012).

### Statistical analyses

Statistical analyses were performed using the Graphpad Prism 6.0 software (Graphpad softwares Inc., La Jolla, USA). Differences in data of two samples were analysed by the student’s t test or ANOVA test with selected comparison using tukey post-hoc test and were considered significant for P-values <0.05 *, p<0.01 ** or p<0.001 ***.

## Results

### Generation and characterization of stable TGF-βRII-KD cellular clones

Expression of TGF-βRI, TGF-βRII, TGF-βRIII Smad2, Smad3, and Smad7 was confirmed by RT-PCR in CAPAN-1 and CAPAN-2 cells. Wild type Smad4, as it is mutated, is not detected in CAPAN-1 (Jonckheere et al., 2004; Schutte et al., 1996). Altogether this suggests that CAPAN-2 cells harbor a functional TGF-β signaling pathway whereas the canonical Smad pathway is not functional in CAPAN-1 cells. Moreover, strong TGF-β1 and mild TGF-β2 mRNA levels were observed in both cell lines suggesting TGF-β growth factor autocrine expression (Figure 1A).

**Figure 1:**
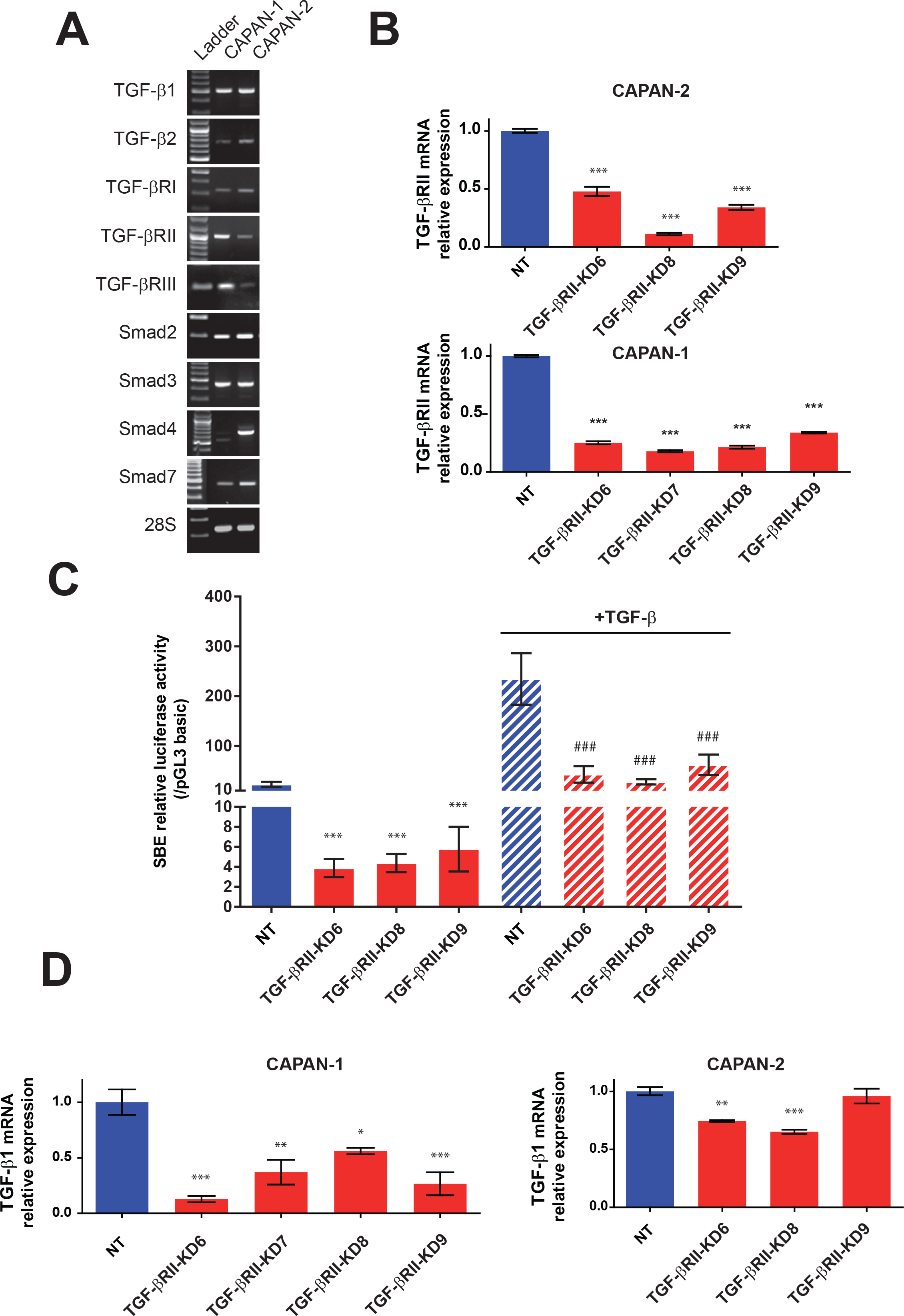
Establishment of TGF-βRII-KD CAPAN-1 and CAPAN-2 cell lines. (A) Analysis of mRNA expression of TGF-β1, TGF-β2, TGF-βRI, TGF-βRII, TGF-βRIII, Smad2, Smad3, Smad4, Smad ATP7A, ATP7B and 28S in CAPAN-1, CAPAN-2 cells by RT-PCR. (B) Analysis of mRNA relative expression of TGF-βRII in NT and TGF-βRII-KD CAPAN-1 and CAPAN-2 cell lines. Expression in NT cells was arbitrarily set to 1. (C) Smad-Binding-Elements (SBE) relative luciferase activity in untreated and TGF-β treated NT and TGF-βRII-KD CAPAN-2 cells. Relative luciferase activity was expressed as a ratio of SBE-Luc normalized with pGL3 basic activity. (D) Analysis of mRNA relative expression of TGF-βi in NT and TGF-βRII-KD CAPAN-1 and CAPAN-2 cell lines.

We generated CAPAN-1 and CAPAN-2 stable cell lines in which TGF-βRII was knocked down (TGF-βRII-KD) by a shRNA approach. Four different shRNA sequences were used to establish four different cell lines designated as TGF-βRIIKD6, TGF-βRIIKD7, TGF-βRIIKD8 and TGF-βRIIKD9. Using qPCR, we confirmed that TGF-βRII mRNA levels are decreased in all CAPAN-1 and CAPAN-2 TGF-βRII-KD cells compared to NT control cells (p:0.005, ***) (Figure 1B). We were not able to produce TGF-βRIIKD7 cell line in CAPAN-2.

In CAPAN-2 KD cells, the inhibition of TGF-βRII expression was correlated with a loss of activity of the Smad binding elements (SBE)-Luc synthetic promoter (Figure 1C). In CAPAN-2 NT cells, TGF-β treatment induces a 10-fold increase of SBE-Luc relative activity whereas this effect was lost in TGF-βRII-KD cells (p<0.001). As expected, in CAPAN-1 cells mutated for Smad4, we did not observe any activity of SBE-Luc construct with or without TGF-β treatment (not shown). Interestingly, TGF-βRII knocking down led to decreased TGF-β1 mRNA level in CAPAN-1 TGF-βRII-KD cells (44-87% decrease) (Figure 1D) whereas the effect was less pronounced (21-25%) in TGF-βRII-KD CAPAN-2 cell lines.

### Involvement of TGF-βRII in PC cell biological properties

We investigated the effect of TGF-βRII silencing on CAPAN-1 and CAPAN-2 proliferation and migration properties. Cell migration was assessed by wound healing test. In CAPAN-2 NT cells, the wound was entirely closed at 60h. In CAPAN-2 TGF-βRII-KD cells, we observed a strong delay of wound closure that was statistically significant at 16-18h (p<0.001, ***) (Figure 2A, left panel). Interestingly, we did not observe any statistically significant difference in wound closure in CAPAN-1 TGF-β-RIIKD or NT cells suggesting the involvement of a functional Smad4 signaling pathway in wound closure (Figure 2A, right panel). TGF-βRII-KD CAPAN-1 or CAPAN-2 cells also showed a trend toward increased proliferation at 96h compared to the respective NT control cells but that remained not significant (not shown).

**Figure 2:**
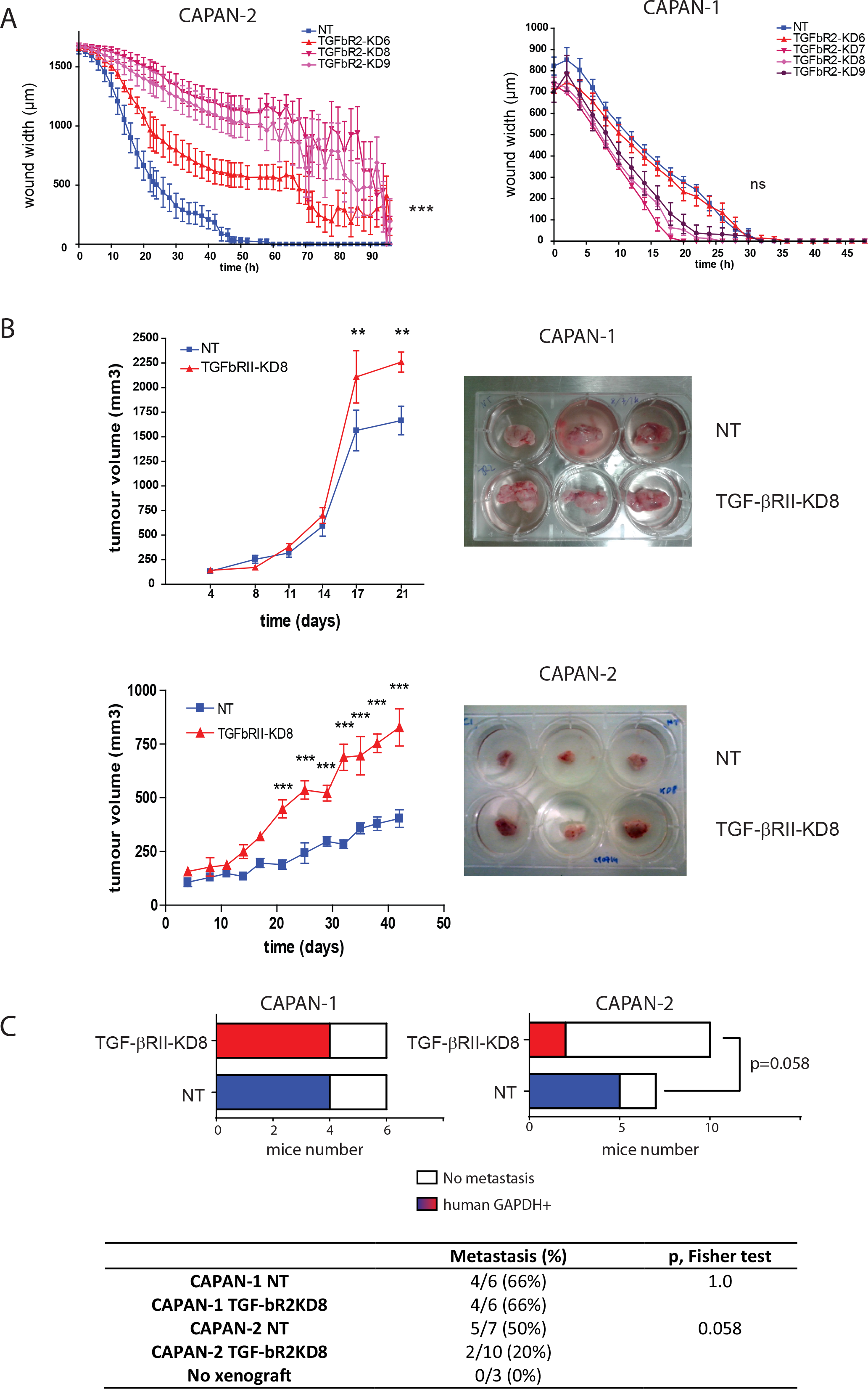
TGF-βRII alters tumor growth and migration in pancreatic cancer cells. (A) Wound healing closure of NT and TGF-βRII-KD CAPAN-1 and CAPAN-2 cell lines using the IncuCyte™ chamber apparatus. (B) Subcutaneous xenografts of NT/TGF-βRII-KD8 CAPAN-1 and CAPAN-2 cells in scid mice. Tumour growth (mm3) was evaluated until sacrifice. **p < 0.01 and ***p < 0.001 indicate statistical significance of TGF-βRII-KD1 compared with the NT control. Ns: not significant. (C) Evaluation of the presence of micro-metastasis in the liver by detecting the presence of human GAPDH in the liver of NT and TGF-βRII-KD CAPAN-1 and CAPAN-2 xenografted mice by qPCR.

In order to determine the role of TGF-βRII on pancreatic carcinogenesis in vivo, CAPAN-1/-2 TGF-βRII-KD8 and NT SC xenograft studies were carried out. We selected the TGF-βRII-KD8 cell lines for in vivo studies as this cell line harboured the best KD in CAPAN-1 and CAPAN-2. The results indicate that the tumour volume was significantly higher in xenografted mice with CAPAN-1 TGF-βRII-KD8 compared to CAPAN-1 NT controls. The relative tumour volume was 2.26±0.1 cm^3^ when compared to NT control tumour volume (1.66±0.14 cm^3^) at day 21. The increase was statistically significant (**, p<0.01). Similar results were obtained with CAPAN-2 TGF-βRII-KD8 xenografts (0.423±0.05 vs 0.828±0.08 cm^3^) at day 42 (Figure 2B). Furthermore, we also evaluated the presence of micro-metastasis in the liver by detecting the presence of human GAPDH in the liver of the mouse by qPCR (Figure 2C). We detected micro-metastases in 5/7 (71%) CAPAN-2 controls whereas only 2/10 (20%) of CAPAN-1 TGF-βRII-KD8 xenografted mice harboured micro-metastases. Contingency analysis showed that difference was close to statistical significance (p=0.058). We did not observe any difference in CAPAN-1 TGF-βRII-KD8 (4/6) compared to CAPAN-1 NT controls (4/6). No human GAPDH mRNA was detected in mice without xenografts. Our results suggest that TGF-βRII signalling is involved in tumor growth and migration of pancreatic cancer cells both in vitro and in vivo.

### Role of TGF-βRII on PC cells sensitivity to gemcitabine

We investigated the effect of TGF-βRII silencing on CAPAN-1 and CAPAN-2 cell sensitivity to gemcitabine. We show that the lack of TGF-βRII induces a significant increase of resistance to gemcitabine treatment in both CAPAN-1 (87-152% increase of survival rate, Figure 3A) and CAPAN-2 (50-161% increase, Figure 3B) cell lines compared to NT control cells. All differences were statistically significant. We then carried out SC xenograft of NT or TGF-βRII-KD8 CAPAN-2 cells that were subsequently treated with gemcitabine for 46 days. Gemcitabine treatment decreased the normalized tumor volume in CAPAN-2 NT xenografts (2.7±1.02 vs 1.4±0.11 at D83) compared to initial tumor volume (D36). On the contrary, the tumor growth was exacerbated in TGF-βRII-KD xenografts following gemcitabine treatment (2.9±1.7 vs 4.36±1 at D83) (Figure 3C). Altogether, our results suggest that TGF-βRII alters sensitivity of PC cells to gemcitabine both in vitro and in vivo.

**Figure 3:**
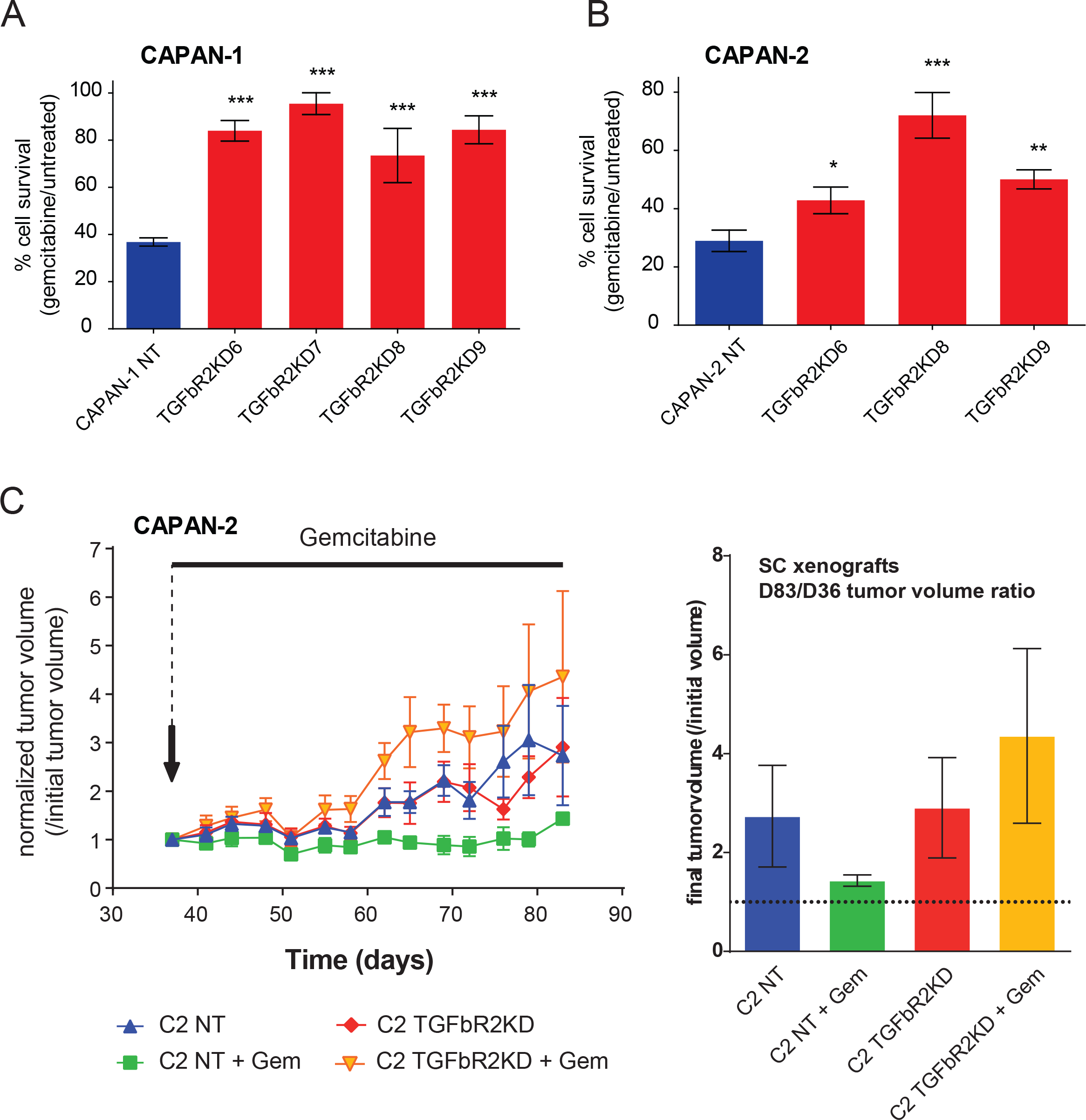
TGF-βRII alters sensitivity to gemcitabine in pancreatic cancer cells in vitro and in vivo. Survival rates in different TGF-βRII-KD CAPAN-1 (A) and CAPAN-2 (B) cell lines or their NT control cells were measured following treatment with gemcitabine using the MTT assay. Results are expressed as % of cell survival (/untreated cells). Three independent experiments were performed. (C) Subcutaneous xenografts of NT and TGF-βRII-KD8 CAPAN-2 cells in scid mice. Gemcitabine (15 mg/kg) or PBS (200 μl) were injected intra-peritoneously, twice a week once palpable tumors were developed. Normalized tumor growth is expressed as the ratio of tumor progression relative to tumor volume on the first day of gemcitabine treatment. Right graph represents tumor growth over time. Left graph represents final tumor volume at day 83 (normalized as initial tumor volume at D36 equal to 1).

### Identification of signalling pathways altered following TGF-βRII knocking down

Impact of TGF-βRII knocking-down on intracellular signaling was studied using phospho array that detect relative site-specific phosphorylation of 43 proteins simultaneously (Figure 4). Intensities of each spots for TGF-βRII were measured and normalized to the CAPAN-2 NT proteins (Figure 4A). We observed an important increase of phosphorylation of S63 c-Jun (3.3-fold) and S727STAT3 (1.5-fold) in CAPAN-2 TGF-βRII-KD8 compared to NT cells. We also observed a mild decrease of phosphorylation of Y694 STAT5a (0.6-fold) and β-catenin (0.5-fold) (Figure 4B). Similar experiments were conducted for CAPAN-1 TGF-βRII-KD8 and NT cells. Only weak variations were observed (30%). By western blotting, we confirmed the increased of phospho-S727 STAT3 (4.49-fold) (Figure 5A) and phospho-S63 (7-fold) (supplemental figure 1A) in CAPAN-2 TGF-βRII-KD8 cells compared to CAPAN-2 NT cells.

**Figure 4:**
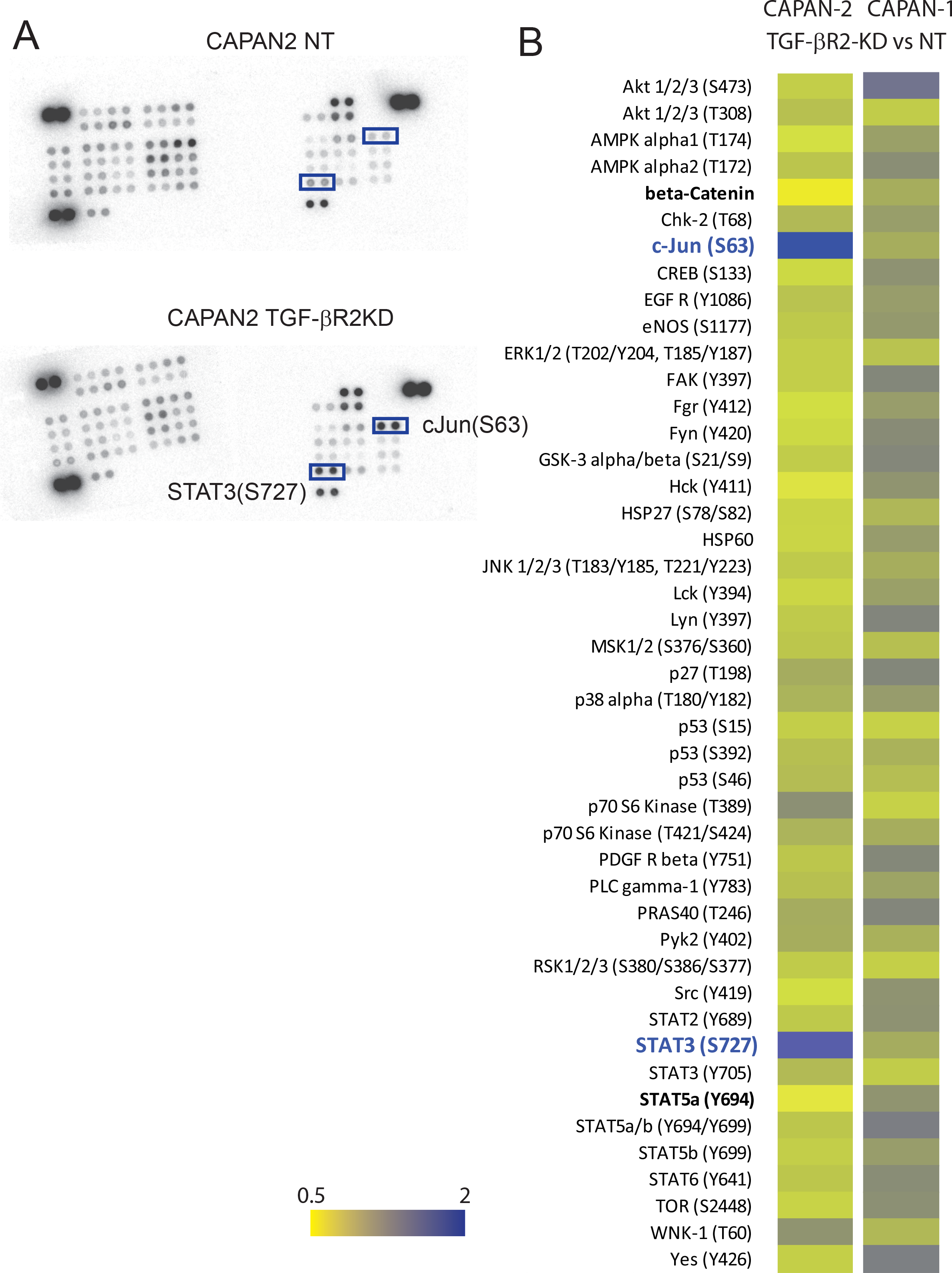
Impact of TGF-βRII knocking-down on signaling pathways. (A) Impact of TGF-pRII knocking-down on intracellular signaling was studied using phospho array that detect relative site-specific phosphorylation of 43 proteins. Boxes highlight spots for S63 c-Jun and S727 STAT3. (B) Heatmap representing the intensities of each spots (TGF-βRII vs NT) that were measured and normalized to the reference spots for CAPAN-1 and CAPAN-2 cells.

**Figure 5:**
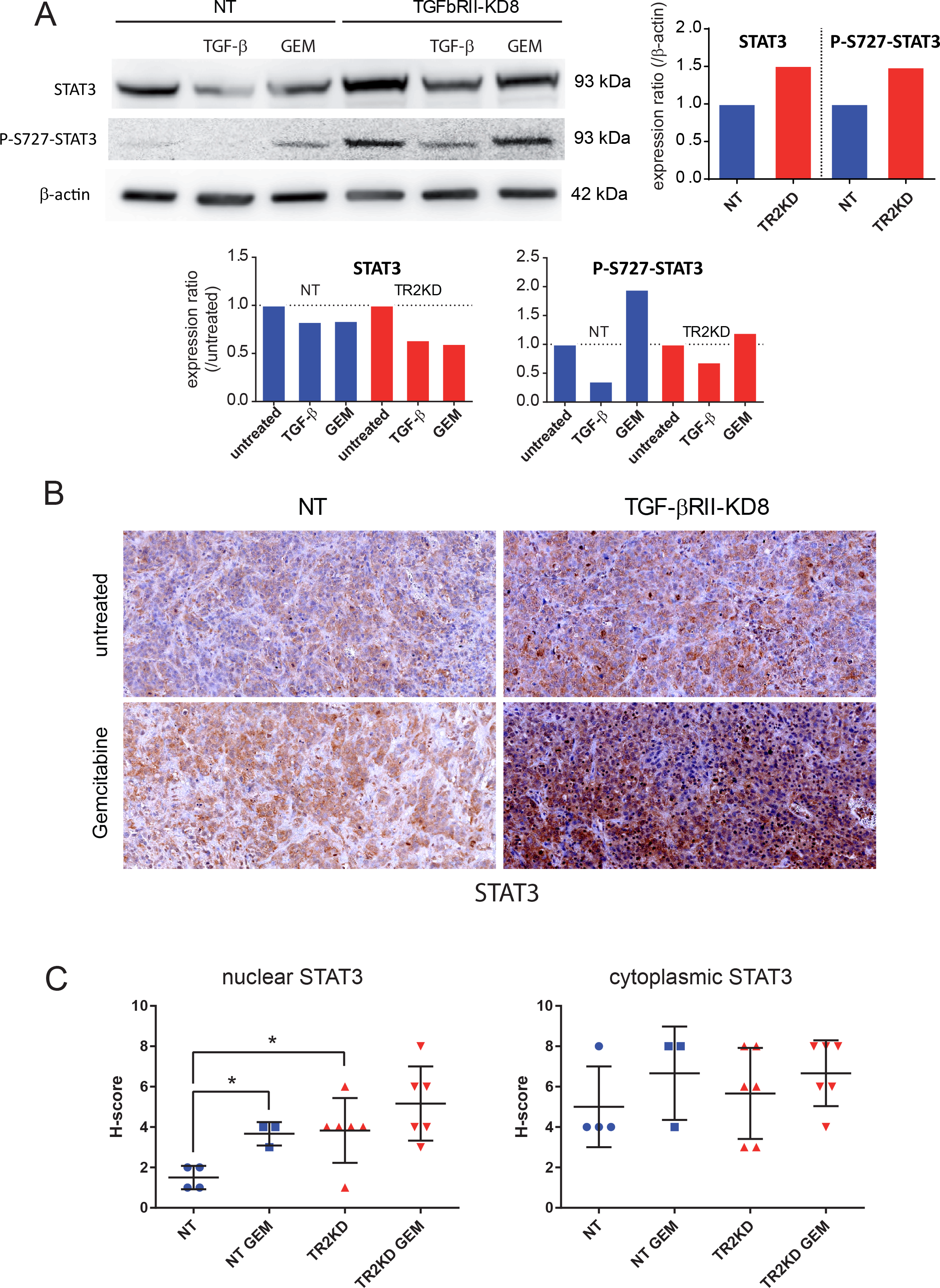
TGF-βRII knockdown promotes STAT3 phosphorylation and nuclear localisation in CAPAN-2 cells. (A) STAT3, phospho-S727 STAT3 and p-actin expression was analysed by western blotting. Bands intensities were quantified by densitometry and ratios (KD vs NT or treated/untreated) are indicated in the graphs. Expression in NT (for TGF-βRIIKD) or untreated (for gemcitabine/TGF-β) cells was arbitrarily set to 1. (B) IHC analysis of STAT3 on extracted xenografted NT and TGF-βRIIKD tumors. (C) Nuclear and cytoplasmic IHC staining were scored in NT and TGF-βRIIKD xenografted tumors that were treated with gemcitabine or PBS. *p<0.05 indicates statistical significance of TGF-βRII-KD1 compared with the NT control.

Gemcitabine treatment also induced an increase of phospho-S727 STAT3 (1.95-fold) (Figure 5A) and phospho-S63 c-Jun (3.18-fold) (supplemental figure 1A) in NT cells (compared to untreated cells). This effect was not found in in TGF-βRII-KD8 cells. We then performed immunohistochemistry for STAT3 and c-Jun in NT or TGF-βRII-KD8 CAPAN-2 SC xenografts (Figure 5B and supplemental figure 1B). Nuclear and cytoplasmic IHC staining were scored. We show that STAT3 nuclear H-score in TGF-βRII-KD8 tumors was significantly higher than in NT tumors (*, p=0.0429) (Figure 5C). We also observed that STAT3 nuclear staining was increased following gemcitabine treatment (*, p=0.0286). A mild increase of nuclear STAT3 was observed in TGF-βRII-KD8 tumors following gemcitabine treatment but was not statistically significant (p=0.33) (Figure 5C). No alteration of c-jun expression was observed in NT and TGF-βRII-KD8 untreated xenograft tumors (supplemental figure 1B). H score measurement indicates that gemcitabine treatment led to a significant decrease of cJun staining in TGF-βRII-KD8 tumors (Supplemental figure 1C). Altogether, our results indicate that TGF-βRII signalling implicates STAT3 and c-Jun phosphorylation in pancreatic cancer cells.

### TGF-βRII silencing alters the expression of ABC transporters and EMT markers in PC cells

To go further and understand which molecular mechanisms could be responsible for the induced chemoresistance, we investigated the effect of TGF-βRII silencing on the expression of ATP-binding cassette (ABC) transporters that are commonly known to confer resistance to xenobiotics including chemotherapeutic drugs. Using qPCR, we investigated the expression of ABCB1/MDR1, ABCC1/MRP1, ABCC2/MRP2, ABCC3/MRP3, ABCC4/MRP4, ABCC5/MRP5 and ABCG2 in NT and TGF-βRII-KD CAPAN-1 and CAPAN-2 cells. MRP1 was not detected. We found that MDR1 (x4.2-fold, **), ABCG2 (x1.9-fold, ***) and MRP4 (x1.4-fold, *) mRNA levels were significantly increased in TGF-βRII-KD CAPAN-1 cells compared to NT cells (Figure 6A). MRP3 mRNA level was decreased in TGF-βRII-KD CAPAN-1 cells (x0.42-fold, ***) and CAPAN-2 (x0.65-fold, p=0.13) (Figure 6A). TGF-βRII and ABC transporter expression was analyzed from 44 pancreatic cancer cell lines from CCLE. We showed that TGF-βRII mRNA relative level was correlated with expression of MRP3 (Pearson r=0.3856, p=0.0097) (Figure 6B) and inversely correlated with MRP4 (Pearson r=−0.3691, p=0.037) (Figure 6C).

Furthermore, TGF-β is commonly described as an inducer of epithelial-mesenchymal transition (EMT) that is associated with chemoresistance (Voulgari & Pintzas, 2009). We performed vimentin (mesenchymal marker) and E-cadherin (epithelial marker) immunohistochemical staining on FFPE sections of NT or TGF-βRII-KD CAPAN-2 xenografts treated with gemcitabine to check their status. Surprisingly, we observed a slight increase of vimentin in TGF-βRII-KD CAPAN-2 cells xenografts compared to NT tumors (supplemental figure 2). Moreover, we found that gemcitabine treatment induced a loss of E-cadherin staining and a gain of vimentin staining suggesting an EMT phenotype following gemcitabine treatment.

**Figure 6:**
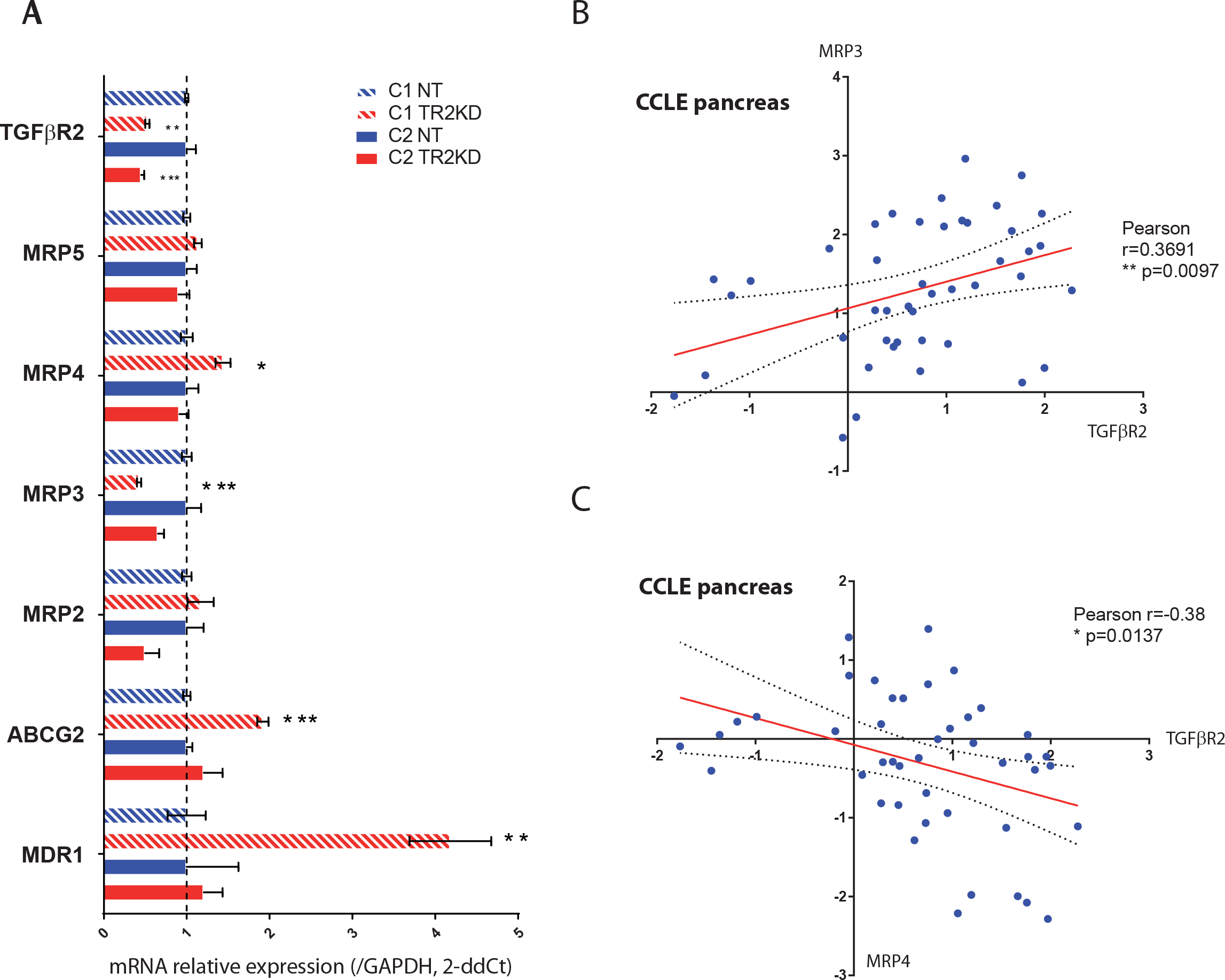
TGF-βRII silencing alters ABC transporters expression. (A) mRNA expression of *TGF-βRII, c-Jun, STAT3, MRP1, MRP2, MRP3, MRP4, MRP5, ABCG2* and *MDR1* was analyzed in NT and TGF-βRII-KD CAPAN-1 and CAPAN-2 cells by qRT-PCR. The histogram represents the ratio of their expression in TGF-βRII-KD compared with NT cells. Three independent experiments were performed. *p<0.05, **p < 0.01 and ***p < 0.001 indicate statistical significance of TGF-βRII-KD1 compared with the NT control. TGF-βRII, MRP3 (B) and MRP4 (C) mRNA expression was extracted from pancreatic cell lines from Cancer Cell Line Encyclopedia (CCLE). Statistical analyses of MRP3/TGFbRII and MRP3/TGF-βRII correlations were analysed Pearson’s correlation coefficient.

Altogether, these results suggest that TGF-βRII silencing alters MRP3 and MRP4 ABC transporters expression in pancreatic cancer cells and induces a partial EMT phenotype that could lead to chemoresistance to gemcitabine.

## Discussion

In the present manuscript, we characterized pancreatic cancer cell lines stably invalidated for TGF-βRII and investigated the consequences on both biological properties and response to chemotherapy in vitro and in vivo. We show an increase of tumor growth and a reduction of cell migration. We also show for the first time an increased resistance to gemcitabine that could be mediated by S727 STAT3 phosphorylation and via deregulation of MRP3 and MRP4 ABC transporter expression. TGF-β signalling pathway has been described as a double edge sword during carcinogenesis (Akhurst & Derynck, 2001); acting as a tumor suppressor in the early stages but promoting metastasis in the advanced carcinoma (Principe et al., 2014). Moses’s laboratory generated TGF-βRII knock out mice crossed with Ptf1a-Cre; LSL-KrasG12D and showed that compound mice developed well differentiated PDAC (Ijichi et al., 2006) mostly highlighting the role as a tumor suppressor. TGFbR2 targeting by a monoclonal antibody is also effective at reducing metastasis (Ostapoff et al., 2014). In our cellular models, we confirmed that TGF-βRII inhibition led to an increased tumor growth in vivo and TGF-βRIIKD CAPAN-2 tumors led to less metastasis in the liver.

STAT3 targeting has been proposed as a therapeutic target in pancreatic adenocarcinoma (Sahu et al., 2017). Combined treatments of gemcitabine and a JAK inhibitor (AZD1480) led to stroma remodeling, increased density of microvessel, enhanced drug delivery and improved survival of in Ptf1a-Cre; LSL-KrasG12D; TGF-βRII^KO^ vivo models suggesting an effect of the treatment via the stroma (Nagathihalli et al., 2015). S727 phosphorylation was previously studied in prostate carcinogenesis and was shown to promote cell survival and cell invasion (Qin et al., 2008). In the present work, we also showed that TGF-βRII inhibition led to STAT3 S727 phosphorylation and increased gemcitabine resistance of the tumor cells suggesting the crucial role of STAT3 in both tumor and stromal cells. STAT3 knockdown was shown to be associated with increased response to gemcitabine in pancreatic cancer cells (Venkatasubbarao et al., 2013). It is interesting to note that Erlotinib treatment that inhibits epidermal growth factor receptor (EGFR) tyrosine kinase also inhibited phosphorylation of STAT3 (Miyabayashi et al., 2013). Among the targeted therapy for PDAC, erlotinib associated with gemcitabine is the only drug showing statistically significantly improved survival (Moore et al., 2007).

TGF-β signaling is mediated through canonical SMAD and non- canonical non-SMAD pathways (Principe et al., 2014). Accordingly, we only observed increase of c-Jun and STAT3 phosphorylation in CAPAN-2 TGF-βRIIKD cells but not in the CAPAN-1 model that is SMAD4 mutated. It was previously shown that STAT3-induced senescence requires functional TGFβR signaling and notably a functional SMAD3/SMAD4 pathway. STAT3 promotes SMAD3 nuclear localization (Bryson et al., 2017). We hypothesize that the TGF-βRIIKD-induced gemcitabine resistance, shown in the present manuscript, is mediated by STAT3 which similarly requires SMAD3/SMAD4 dependent pathway.

In Smad4 mutated CAPAN-1 cells we observed an increased expression of MRP4, ABCG2 and MDR1. This increased expression could be responsible for the gemcitabine resistance of CAPAN-1 TGF-βRIIKD cells. The link between ABC transporters and TGF-β pathway is scarcely described. TGF-β1 has been shown to upregulate ABCG2 expression in MiaPACA2 pancreatic cells which is contradictory with our findings (Kali et al., 2017). In breast cancer cells, silencing of TGF-βRII leads to overexpression of multidrug resistance protein ABCG2 and tamoxifen resistance (Busch et al., 2015).

TGF-β is usually considered as a bona fide inducer of EMT (Wendt et al., 2009). However, we were surprised to observe that TGF-βRII inhibition led to a partial EMT with an increase of vimentin. STAT3 signaling is linked to cancer cell plasticity and is able to promote EMT and CSC expansion (Junk et al., 2017). Previous work also showed that IL6, secreted by pancreatic stellate cells, triggers STAT3 activation in pancreatic cells which subsequently induces EMT via Nrf2 (Wu et al., 2017). Therefore, we hypothesize that the paradoxal EMT observed in TGF-βRII cells is a consequence of the STAT3 phosphorylation on S727.

## Aknowledgments

Vincent Drubay is a recipient of a fellowship “Année recherche” from the Lille University/Faculty of Medicine. Nicolas Skrypek is a recipient of a PhD fellowship from the Centre Hospitalier Régional et Universitaire (CHRU) de Lille/région Nord-Pas de Calais. Romain Vasseur and Nihad Boukrout are recipients of a PhD fellowship from the Université of Lille 2. Isabelle Van Seuningen is the recipient of a “Contrat Hospitalier de Recherche Translationnelle”/CHRT 2010, AVIESAN. This work is supported by Inserm and CNRS and grant from la Ligue Nationale Contre le Cancer (Equipe Labellisée Ligue 2010, IVS; Ligue comité 62, NJ) and by SIRIC ONCOLille, Grant INCaDGOS-Inserm 6041 and by a grant from “Contrat de Plan Etat Région” CPER Cancer 2007-2013. We thank MH Gevaert (Department of Histology, Faculty of Medicine, University of Lille) for their technical help and the University of Lille EOPS animal facility (D. Taillieu).

## Supplemental material

**Supplemental Figure 1: TGF-βRII knockdown promotes c-Jun S-63 phosphorylation in CAPAN-2 cells.** (A) c-Jun, phospho-S63 c-Jun and β-actin expression was analysed by western blotting. Bands intensities were quantified by densitometry and ratios (KD vs NT or treated/untreated) are indicated in the graphs. Expression in NT (for TGF-βRIIKD) or untreated (for gemcitabine/TGF-β) cells was arbitrarily set to 1. (B) IHC analysis of c-Jun on extracted xenografted NT and TGF-βRIIKD tumors. (C) IHC staining was scored in NT and TGF-βRIIKD xenografted tumors that were treated with gemcitabine or PBS. *p<0.05 indicate statistical significance of TGF-βRII-KD compared with the NT control.

**Supplemental Figure 2: TGF-βRII knockdown promotes partial EMT-like phenotype.** IHC analysis of E-cadherin and vimentin on extracted xenografted NT and TGF-βRIIKD tumors. IHC staining was scored in NT and TGF-βRIIKD xenografted tumors that were treated with gemcitabine or PBS. *p<0.05, **p<0.01 indicate statistical significance of TGF-βRII-KD compared with the NT control.

